# Real-space refinement in *Phenix* for cryo-EM and crystallography

**DOI:** 10.1101/249607

**Authors:** Pavel V. Afonine, Billy K. Poon, Randy J. Read, Oleg V. Sobolev, Thomas C. Terwilliger, Alexandre Urzhumtsev, Paul D. Adams

## Abstract

This article describes the implementation of real-space refinement in the *phenix.real_space_refine* program from the *Phenix* suite. Use of a simplified refinement target function enables fast calculation, which in turn makes it possible to identify optimal data-restraints weight as part of routine refinements with little runtime cost. Refinement of atomic models against low-resolution data benefits from the inclusion of as much additional information as is available. In addition to standard restraints on covalent geometry, *phenix.real_space_refine* makes use of extra information such as secondary-structure and rotamer-specific restraints, as well as restraints or constraints on internal molecular symmetry. Re-refinement of 385 cryo-EM derived models available in the PDB at resolutions of 6 Å or better shows significant improvement of models and the fit of these models to the target maps.

**Synopsis:** A description of the implementation of real-space refinement in the *phenix.real_space_refine* program from the *Phenix* suite and its application to re-refinement of cryo-EM derived models.

## 1. Introduction

Improvements in the electron cryo-microscopy technique (cryo-EM) have led to a rapid increase in the number of high-resolution three-dimensional reconstructions that can be interpreted with atomic models (Figure 1). This has prompted a number of new developments in *Phenix* (Adams *et al*, 2010) to support the method, from model building (Terwilliger *et al.*, 2018, submitted), map improvement (Terwilliger, manuscript in preparation) and refinement (Afonine *et al.*, 2013) to final model validation (Afonine *et al.*, 2018). In this manuscript we focus on atomic model refinement using a map (primarily cryo-EM but the same algorithms and software are also applicable to crystallographic maps).

**Figure 1.**
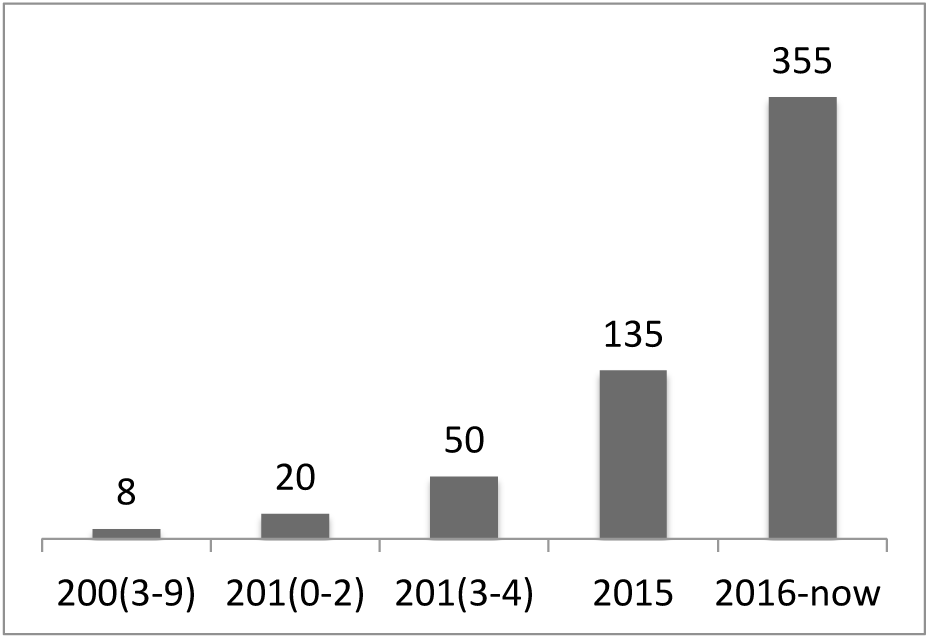
Number of cryo-EM derived models in PDB at resolutions of 6Å or better.

Model refinement is an optimization problem and as such it requires defining three entities (for reviews, see Tronrud, 2004; Watkin, 2008; Afonine *et al.*, 2012, 2015). *The model, i.e.* a mathematical construct that explains the experimental data, with an associated set of refinable parameters – in this case an atomic model with coordinates whose positions can be varied to improve the fit to the data. *The target function* that links the model parameters to the experimental data - this function scores model to data fit, and therefore guides refinement. Finally, *an optimization method* that changes the values of refinable model parameters such that the model agreement with the experimental data is improved. If the target function is expressed through diffraction intensities or structure factors, refinement is usually referred to as reciprocal-space, or Fourier-space refinement (FSR). Alternatively, a target function may be formulated in terms of a map, a Fourier synthesis in the case of crystallography or a three-dimensional reconstruction from projections in the case of cryo-EM. Such refinement is referred to as real-space refinement (RSR). In both cases the targets are the sums over a large number of similar terms corresponding to either reflections (FSR) or map grid points (RSR). A key methodological difference is that for RSR each term depends on only a few atoms, while for FSR each term depends on all model parameters. Most modern macromolecular refinement programs were developed for crystallographic data and therefore perform refinement in reciprocal space, at least as their main mode of operation (see Table 1 in Afonine *et al*, 2015). This work focuses on the realspace refinement of coordinates of atomic models.

**Table 1:** Summary of statistics for original (3j5p), re-refined by Barad *et al*. (2015; 3j9j) and rerefined by *phenix.real_space_refine* models.

| Metric | 3j5p<sup>(*)</sup> | 3j9j | 3j5p<sup>(*)</sup><br>(*phenix.real_space_refine*) |
| --- | --- | --- | --- |
| CC<sub>mask</sub> | 0.6494 | 0.5856 | 0.8171 |
| EMringer score | 1.17 | 2.57 | 3.28 |
| RMSD |  |  |  |
| bonds (Å) | 0.008 | 0.015 | 0.010 |
| angles (°) | 1.495 | 1.085 | 1.440 |
| Ramachandran plot (%) |  |  |  |
| favored | 95.77 | 94.49 | 93.32 |
| allowed | 4.23 | 3.27 | 6.68 |
| outliers | 0 | 2.24 | 0 |
| Rotamer outliers (%) | 32.45 | 0 | 0 |
| Clashscore | 100.83 | 2.70 | 5.62 |
| C<sub>β</sub> deviations | 0 | 0 | 0 |

(\*) No ankyrin domain

In cryo-EM studies real-space refinement is a natural choice because a threedimensional map is the output of the single particle image reconstruction method (Frank, 2006) and does not change in a fundamental way as the atomic model is improved. This is not the case for crystallography, where the experimental data are diffraction intensities, and the associated and vital phase information has to be obtained indirectly. In crystallography, obtaining the best phases typically involves their calculation from atomic models, in turn making the resulting maps model-biased (see for example Hodel *et al.*, 1992). Although FSR methods are predominant in crystallographic refinement, RSR is attractive in some contexts as it makes it possible to refine parts of the model locally.

In the case of cryo-EM an atomic model can also be refined using a reciprocal space target. This can be achieved by converting the map into Fourier coefficients. These Fourier coefficients can then be used in reciprocal-space refinement using standard refinement protocols that are well established for crystallographic structure refinement (see for example Cheng *et al.*, 2011; Baker *et al.*, 2013; Brown *et al.*, 2015). We note, however, that unless the map is converted to the full corresponding set of Fourier coefficients (and not a subset containing only a sphere limited to the stated resolution) this conversion may not be lossless.

To address the emerging structure refinement needs of the rapidly growing field of cryo-EM the *phenix.real_space_refine* program (Afonine *et al.*, 2013), capable of refinement of atomic models against maps, has been introduced into the Phenix suite (Adams *et al.*, 2010). It is not limited to cryo-EM and can be used in crystallographic refinement as well (X-ray, electron or neutron). In this paper we describe the implementation of the *phenix.real_space_refine* program, and demonstrate its performance by applications to simulated data and to cryo-EM models in the PDB (Bernstein *et al.*, 1977; Berman *et al.*, 2000) and corresponding maps in the EMDB (Henrick *et al.*, 2003). This is a work in progress, and further details and advances will be reported as the program evolves. To date *phenix.real_space_refine* has been used in a number of documented structural studies (see for example: Fischer *et al.* (2015); Shalev-Benami *et al.* (2016); Chua *et al.* (2016); Ahmed *et al.* (2016); Yang *et al.* (2016); Gao *et al.* (2016); Chen *et al.* (2016); Bhardwaj *et al.* (2016); Lokareddy *et al.* (2017); Hryc *et al.* (2017); Ahmed *et al.* (2017); Demo *et al.* (2017); Paulino *et al.* (2017); Liu *et al.* (2017)).

## 2. Methods

### 2.1 Refinement flow chart

Figure 2 shows the model refinement flowchart as it is implemented in *phenix.real_space_refine*. This is very similar to the reciprocal-space refinement workflow implemented in *phenix.refine* (see Figure 1 in Afonine *et al.*, 2012).

**Figure 2.**
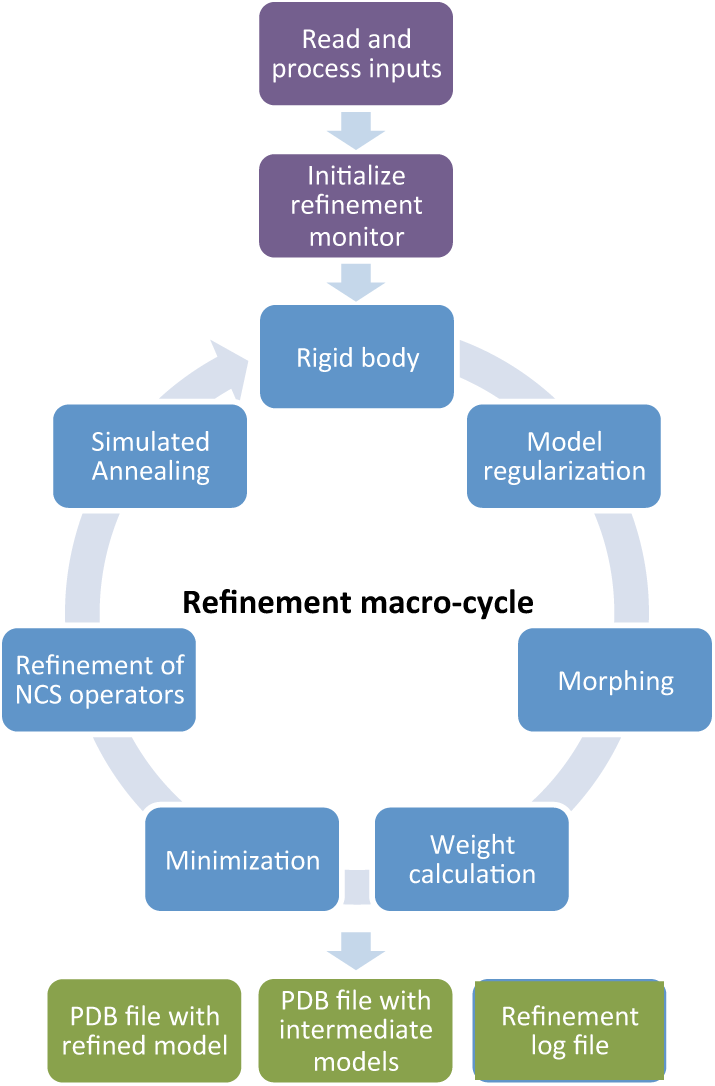
*phenix.real_space_reflne* flowchart.

The program begins by reading a model file, in PDB or mmCIF format, map data (as an actual map in MRC/CCP4 format or as Fourier map coefficients in MTZ format) and other parameters, such as resolution (if a map is provided) or additional restraint definitions for novel ligands, internal molecular symmetry (e.g. NCS in crystallography) or secondary structure. Once inputs are read, the program proceeds to calculations that constitute a set of tasks repeated multiple times (macro-cycles). Tasks to be performed during the refinement are defined by the program automatically and/or by the user. In its default mode the program will only perform gradient-driven minimization of the entire model. Other non-default tasks allow optimization using simulated annealing (SA; Brunger *et al.*, 1987), morphing (Terwilliger *at al.*, 2013), rigid-body refinement (see Afonine *et al.*, 2009 and references within) and systematic residue side-chain optimizations using grid searches in torsion χ-angle space (Oldfield, 2001). Parts of the model related by internal symmetry are determined automatically, if available, or can be defined by the user. In the presence of such internal symmetry, restraints or constraints can be applied between the coordinates of related molecules. The operators relating molecules can be refined as well. The result of refinement, i.e. the refined model, is output as a file in PDB or mmCIF format.

Central to almost all tasks performed within a refinement macro-cycle is the target function. Its choice is the key for the success of refinement, i.e. efficient convergence to an improved model. Also, of the same importance is the assessment of refinement progress by quantifying model quality and the goodness of model-to-map fit throughout the entire process. Some relevant points are discussed below.

### 2.2. Refinement target function

Macromolecular cryo-EM or crystallographic experimental data are almost always of insufficient quality to refine parameters of atomic models individually. To make refinement practical, restraints or constraints are almost always used in order to incorporate extra information into refinement, and the corresponding procedures are called restrained or constrained refinement. In restrained refinement the target function is a sum of data-based and restraints-based components:

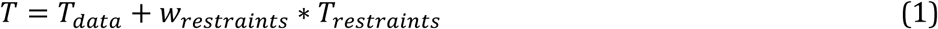

The first term scores the model to data fit and the second term incorporates *a priori* information about the model. The weight *w_restraints_* balances the contribution of restraints to maximize the model-to-data fit while also obeying the *a priori* information. Constrained refinement does not change the target function but rather changes (reduces) the set of independent parameters that can vary. Examples are rigid-body refinement or use of a riding model (Sheldrick & Schneider, 1997) to parameterize the positions of hydrogen atoms.

#### 2.2.1. Model-to-map target (T_data_)

In RSR, the *T_data_* term scores the fit of the model being refined to a target map. In cryo-EM the map is a 3D reconstruction, while in crystallography it may be, for example, a 2*mF_obs_ - DF_model_* map (Read, 1986).

It is possible to express the difference between the two maps in the integral form (e.g. Diamond, 1971)^1^

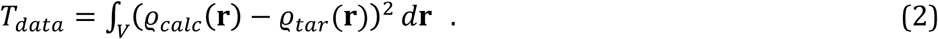

For (2) we suppose that the original target map is optimally scaled to the model map (Diamond, 1971; Chapman 1995). In the following, we will consider the target to be essentially unchanged by manipulations that shift its value by a constant or a scale factor, as such manipulations do not change the position of the minimum of the target. If the Euclidean norms of ϱ_*tar*_**(r)** and ϱ_*calc*_**(r)** are conserved during refinement (*i.e.* if 
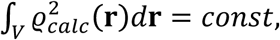
, which will be true if the overlap of atomic densities does not change, and if 
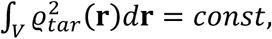
, as will be the case when the target map itself does not change) then minimization of (2) is equivalent to minimization of the anti-correlation target, which does not need the maps to be optimally scaled,

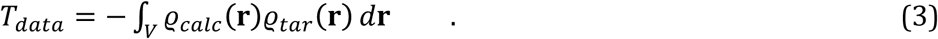

Assuming the target ϱ_*tar*_ and model-calculated ϱ_*calc*_ maps are provided on the same grid, a continuous integration in (2) and (3) can be replaced with a numeric integration over the regular grid on which the maps are available (see, for example, Diamond, 1971),

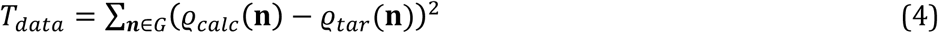

or

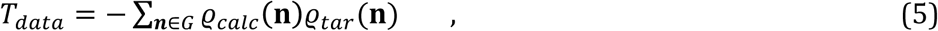

respectively. The set *G* of grid nodes used to calculate the targets (*i.e.* the integration volume) is either the whole map or an envelope (mask) surrounding the atomic model.

To match the finite resolution of the target map in (5) accurately, several steps are required to compute the model map. First, the model density distribution is calculated using one of the available approximations (Neutron News, 1992; Maslen *et al.*, 1992; Waasmaier & Kirfel, 1995; Grosse-Kunstleve, Sauter *et al.*, 2004; Peng *et al.*, 1996; Peng, 1998). Then a set of Fourier coefficients is calculated from the distribution up to the resolution limit specified by the target map^2^. Finally, a subset of these coefficients is used to calculate the model Fourier synthesis ϱ_*calc*_ that can then be used in (5). This synthesis is a representation of a model image at a given resolution. A typical refinement may require hundreds or even thousands of such model image calculations, which are computationally expensive (involving two Fourier transforms).

Alternatively, a model map may be calculated from the atomic model directly as a sum of individual contributions of *M* atoms with each contribution being a Fourier image (or its approximation) of the corresponding atom at a given resolution (e.g. Diamond, 1971; Lunin & Urzhumtsev, 1984; Chapman, 1995; Mooij *et al.*, 2006; Sorzano *et al.*, 2015). While this is much faster than the previous method, it may be less accurate and still be computationally expensive, especially for large models.

A numeric integration over the whole map (5) can be simplified by the integration exploring the volume directly around the atomic centers **r**_*m*_, *m = 1,…M*:

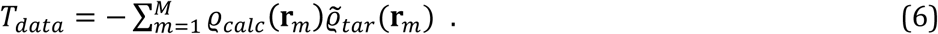

Here 
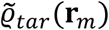
 are the values interpolated from the nearby grid node values ϱ_*tar*_ **(n)** to the atomic centers **r**_*m*_ (Appendices A and B). Neglecting the local variation of the model map at atomic centers (e.g. at low resolution) and thus supposing 
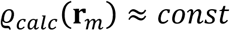
 for all *m*, the target simplifies further as (Rossmann, 2000; Rossmann *et al.*, 2001)

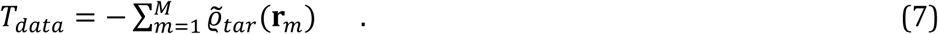

The hypothesis 
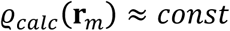
 seems to be reasonable at low resolution when a calculated map can be considered to be rather flat. On the other hand, minimization of (7) is essentially a fitting of atoms to the nearest peaks of the target map, which seems to be appropriate at high resolution as well. We show below (§3) that indeed this target function is efficient over a large resolution range; Appendix B supports this observation through the equivalence of targets (7) and (5) when taking into account map blurring / sharpening. If the difference in atomic size cannot be neglected, this target function can be modified to

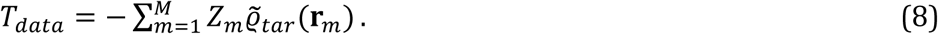

where *Z_m_* is the electron number of the corresponding atom (see also Mooij *et al.*, 2006). Clearly, for most of the macromolecular structures under consideration here these atomcentered targets are nearly the same, and for simplicity in what follows we refer only to (7) unless otherwise stated. The computational cost of (7) is proportional, with a very small coefficient, to the number of atoms and therefore these targets are much faster to calculate compared to (5) making it advantageous for refinement of large models. The *phenix.real_space_refine* program uses it as the default refinement target.

#### 2.2.2. Restraints (T_restraints_)

In restrained refinement, extra information is introduced through the term *T_restraints_* with some weight (1). This extra term restrains model parameters to be similar (but not necessarily identical) to some reference values. At high to medium resolutions, approximately 3 Å or better, a standard set of restraints (Grosse-Kunstleve & Adams; 2004) includes restraints on covalent bond lengths and angles, dihedral angles, planarity and chirality restraints, and a non-bonded repulsion term. However, at lower resolutions the amount of experimental data is insufficient to preserve geometry characteristics of a higher level of structural organization (such as secondary structure) and therefore other information can be included to help produce a chemically meaningful model (Headd *et al*., 2012; Headd *et al*., 2014). These extra restraints may include restraints on similarity of related copies (NCS in the case of crystallography), restraints on secondary structure, and restraints to one or more external reference models (Headd *et al*., 2012, 2014). *phenix.real_space_refine* can use the following extra restraints:

- Distance and angle restraints on hydrogen bond patterns for protein helices and sheets (Headd *et al*., 2012), and DNA/RNA base-pairs;
- Torsion angle restraints on idealized protein secondary structure fragments;
- Restraints to maintain stacking bases in RNA/DNA parallel (parallelity restraints; Sobolev *et al*., 2015);
- Ramachandran plot restraint (Headd *et al*., 2012);
- Amino-acid side-chain rotamer-specific restraints;
- C_β_ deviation restraints;
- Reference model restraints, where a reference model may be a similar structure of better quality or the initial position of the model being refined (Headd *et al*., 2012).
- Similarity restraints in torsion or Cartesian spaces (Headd *et al*., 2014).

#### 2.2.3. Relative weight

The relative weight *w_restraints_* is chosen such that the model fits the map as well as possible while maintaining reasonable deviations from ideal covalent bond lengths and angles. The choice of *w_restraints_* is determined in RSR by systematically trying a range of plausible values and performing a short refinement for each trial value. A similar procedure in FSR would be very computationally expensive because for each trial value of *w_restraints_* the whole structure would need to be used. Fortunately, in RSR this can be performed much faster. The weight calculation procedure implemented in *phenix.real_space_refine* splits the model into a set of randomly chosen segments, each one a few residues long. After trial refinements of each segment with different weights, the best weight is defined as the one that results in a model possessing reasonable bond and angle root-mean-square deviations (rmsd) and that has the best model-to-map fit among all trial weights. The obtained array of best weights for all fragments is filtered for outliers and the average weight is calculated and defined as the best weight for the final refinement. This calculation typically takes less than a minute on an ordinary computer and is independent of the size of the structure or map. Instead of computing an average single weight for the entire model this protocol can in principle be extended (work in progress) to calculate and use different weights for different parts of the map accounting for variations in local map quality.

### 2.3 Evaluation o f refinement progress and results

It is recognized that model validation (for example Branden & Jones, 1990; Read *et al*., 2011; Wlodawer & Dauter, 2017) is a critical step in structure determination and a number of corresponding tools have been developed in crystallography (e.g. Chen *et al*., 2010; Gore *et al*., 2012; Young *et al*., 2017, and references therein) and some in cryo-EM studies (see for example, Henderson *et al*., 2012; Tickle, 2012; Barad *et al*., 2015; Pintilie *et al*., 2016; Joseph *et al*., 2017, Afonine *et al*., 2018). Generally, the process consists of assessing data, model quality and model-to-data fit quality, and is performed locally and globally. At the stage of refining a model we assume that the intrinsic data quality has already been evaluated, and only model quality and model-to-data fit need to be monitored.

Methods and tools to evaluate the geometric quality of a model are the same in crystallography and in cryo-EM. For example, the *Phenix* comprehensive validation program provides an extensive report on model quality, making extensive use of the MolProbity validation algorithms (Chen *et al*., 2010). In crystallography, the model-to-data fit is quantified by crystallographic *R* and *R*_free_ (Brunger, 1992) factors, which are global reciprocal space metrics. In cryo-EM model and data validation is currently performed by the comparison of complex Fourier coefficients in resolution shells; these coefficients are calculated from the model and from the full map or half-maps; different masks can be applied prior to calculation of these coefficients. Also in real-space the model-to-data fit can be evaluated locally or globally by various correlation coefficients between a modelcalculated map and the experimentally derived map. Some of these tools are used in § 3.2 where models extracted from the PDB are refined against experimental cryo-EM maps.

## 3. Results

### 3.1. Test refinements with simulated data

Calculation of (7) requires map interpolation at atomic centers (see Appendix B). Furthermore, refinement requires calculation of the gradients of (7) with respect to atomic coordinates. Calculation of the gradients can involve finite difference approaches (Faddeev & Faddeeva, 1963) or algorithmic derivation (Baur & Strassen, 1983; Kim *et al*., 1984). Here we focus on the three following approaches: linear interpolation with the corresponding gradients (LI), quadratic interpolation with the corresponding gradients (QI) and linear interpolation where the gradients are calculated using finite differences combined with interpolation from neighboring intervals (ID, Interpolated Difference). Other map parameters such as sampling rates, sharpening or blurring can also affect refinement performance. To study the effects of these choices on refinement several numerical tests using simulated data were performed as described below. All refinements were performed using geometry restraints included with optimal weights.

#### 3.1.1. Preparing simulated data

A lysozyme model from the PDB (PDB code 3VB1) was chosen as a test model. The following manipulations were made to this model prior to test calculations: a) the model was placed in a sufficiently large P1 unit cell, b) all atomic displacement parameters (Bfactors) were set to 20 Å^2^, c) alternative conformations were replaced with a single conformation, d) model geometry was regularized using the *phenix.geometry_minimization* tool until convergence. In what follows we refer to this model as a *reference model*. Several Fourier maps at different resolutions *d_high_* (1, 2, 3, 4, 5 and 6 Å) were calculated from the reference model to mimic ϱ_*tar*_. These maps were calculated on a grid with the step equal to *d_high_*/4 (in what follows referred to as a resolution-based grid). Additionally, we calculated the same maps on a much finer grid with a step of 0.2 Å, the same for all maps independent of their resolution.

#### 3.1.2. Refinement o f the exact reference model

First, we refined the reference model against twelve finite resolution maps calculated from this model as described in §3.1.1. While the reference model corresponds to the minimum of (5) this is not the case for (7) because map peaks in finite resolution Fourier images do not necessarily correspond to atomic centers. Therefore, it is expected that refinement using (7) may shift the model from its original, correct, position. The goal of this test is to provide an estimate of the magnitude of these shifts after refinement. Each refinement was repeated three times testing each of the three considered options for map interpolation and gradient calculation. This resulted in 36 refined models (6 map resolutions * 2 sampling rate choices * 3 map interpolation and gradient calculation options). For each refined model we calculated root-mean-square deviation (rmsd) from the reference model. Figure 3 summarizes the result of this test. We observe:

- Independently of map sampling rate, the rmsd increases almost linearly as resolution worsens and ranges from as low as 0.02 Å at 1 Å resolution to as high as 0.43 Å at 6 Å resolution. These rmsds are small compared to the details that can be resolved in maps of these resolutions. This justifies the use of a target (7) that is less accurate but much faster to calculate than (5).
- Refinements using coarser maps show, that

- LI produces the worst results across all resolutions but 6 Å;
- QI produces systematically the best results at resolutions 4 ‒ 6 Å, which can be explained by the absence of sharper map details when a quadratic approximation becomes more accurate than a linear one;
- ID performs best at resolutions 1‒3 Å. This possibly can be explained because gradient calculations involve interpolation from neighboring intervals and thus ID uses more information than LI or QI.

Additionally, we note that relative differences among LI, QI and ID results are small compared to the errors introduced by any of the methods.

- #x002D; Refinement using finely-sampled maps does not significantly change the rmsd obtained with ID compared to refinement using coarsely-sampled maps. This shows that these rmsd values are not due to a finite grid step but are an intrinsic limitation of the target (7) that does not account for displacement of the map peaks from the atomic centers. Also, we note that, as expected, using finely-sampled maps almost eliminates the differences between LI, QI and ID.

**Figure 3.**
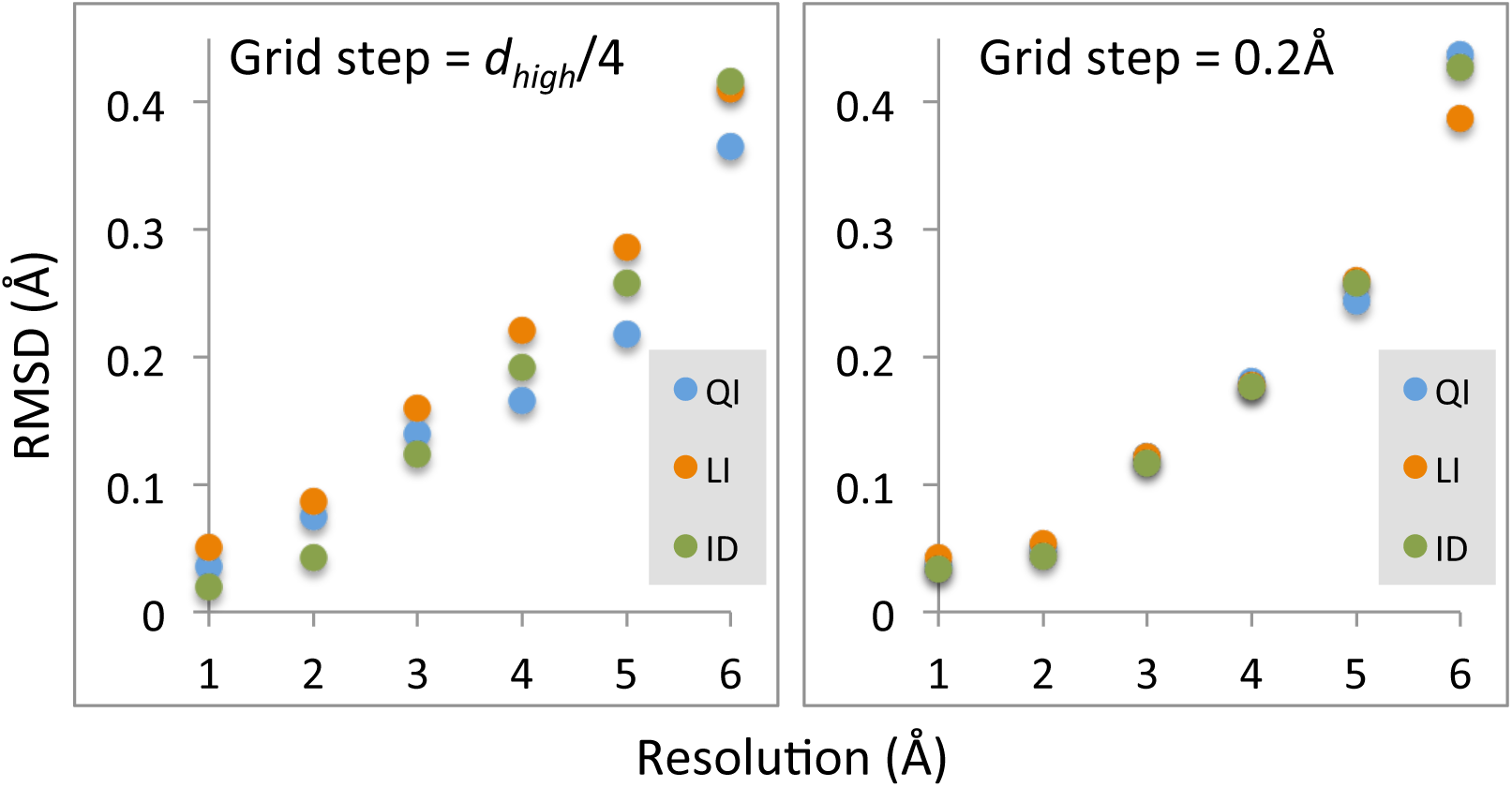
Refinement of the exact model against six maps computed on the resolution-dependent grid (left) and six maps computed on the finer fixed grid (right). Each circle shows root-mean-square deviation between refined model and the reference model, for each of twelve maps and for each of interpolation/gradient calculation methods (QI, LI and ID). See §3.1.2 for details.

Next, we tested how map sharpness could influence the refinement. Intuitively, it seems likely that it may be difficult for refinement to find the correct atomic positions if peaks are broad and so several atoms are close to the same peak (in the case of a blurred map). On the other hand, if peaks are too sharp, atoms may not be within the radius of convergence, which is problematic for refinement as well. To illustrate the point, we used six maps from §3.1.1 of resolutions from 1 to 6 Å calculated on the grid with the step of *d_high_*/4 and modified these maps to correspond to the reference model having B-factors from 0 to 200 Å^2^. Then we refined the reference model against each of these maps.

Figure 4a shows the rmsd between refined and reference models as a function of overall B-factor for six resolutions. At 1-2 Å resolution the exact maps calculated with B=0 Å2 show sharp peaks for atomic positions so any map blurring makes refinement worse. At 6 Å resolution the map with B=0 Å2 is already too featureless to contain atomic information and any additional blurring seems to be counter-productive. On the other hand, sharpening the map in this case isn’t helpful either (Figure 4b) because it does not recover individual atomic peaks. At intermediate resolutions, from 3 to 5 Å, neither very sharp (B=0 Å2) nor very blurred (B=200 Å2) maps are the best choice. A possible explanation for this may be the following. At these resolutions the peaks are still strong (compared to 6 Å resolution maps) but they are not centered on atomic positions. Minimization of (7) may push atoms into these peaks, which isn’t desirable. Therefore, some attenuation of these peaks may be useful, as Figure 4b illustrates.

**Figure 4.**
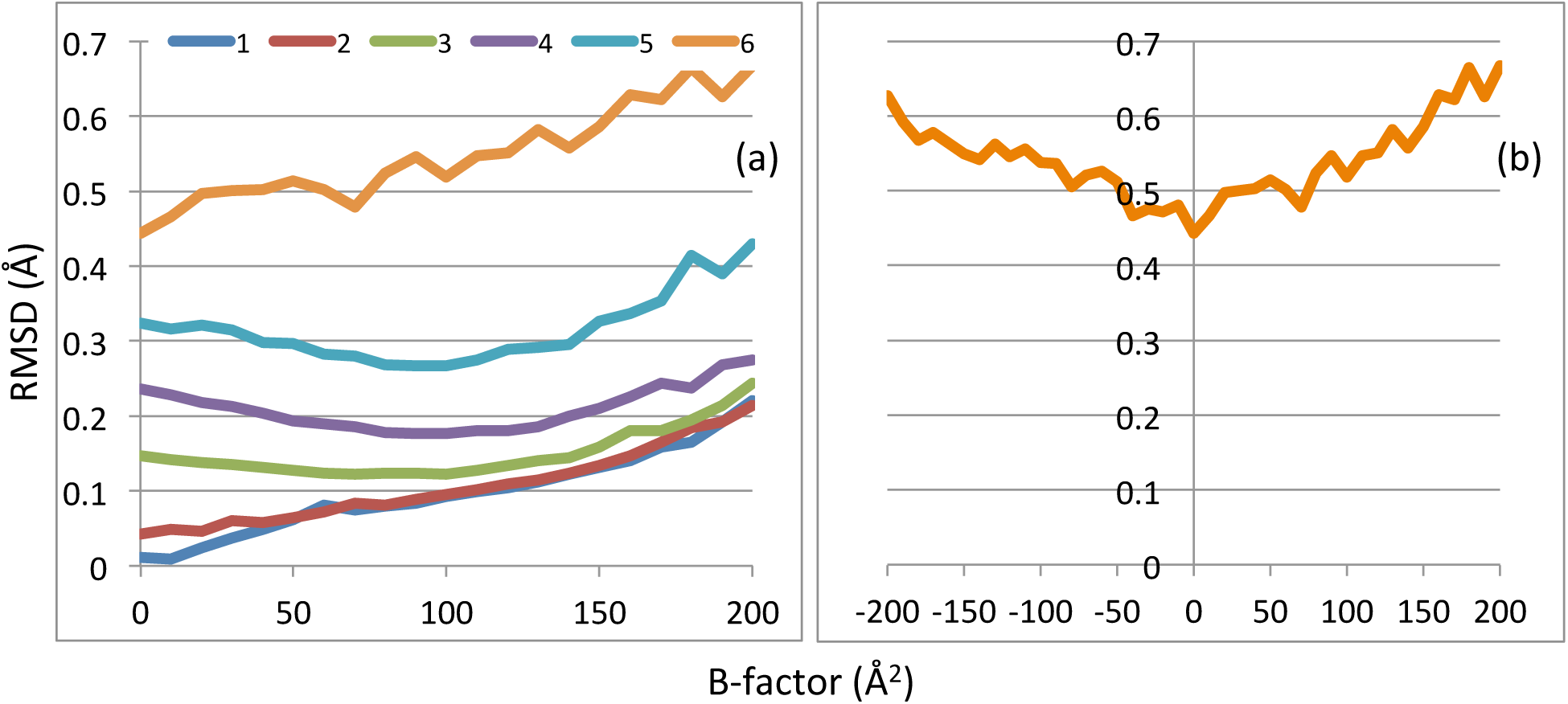
Effect of map sharpness. RMSD between reference and refined models at resolutions (a) from 1 to 6Å and (b) 6Å, shown as function of overall B-factor. See text for details.

#### 3.1.3. Refinement o f perturbed reference models

Here we describe tests that are similar to those from §3.1.2 except that instead of refining the reference model we refined perturbed reference models. The perturbed models were obtained by running molecular dynamics (MD) simulations using the *phenix.dynamics* tool until a prescribed rmsd compared to the reference model was achieved. Given the stochastic nature of MD it is possible to obtain many different models with the same rmsd from the reference model. Due to the limited convergence radius of refinement and finite resolution of the data, refinement of these models will not produce exactly the same refined models. Therefore, to ensure more robust statistics, for each chosen rmsd we generated an ensemble of 100 models. The rmsd values between perturbed and reference models were chosen to be 0.5, 1.0, 1.5 and 2.0 Å. Then we refined each of these 100 × 4 = 400 models against six maps (at resolutions *d_high_* from 1 to 6 Å) calculated on a grid with the spacing *d_high_*/4. Similarly to the previous test, each refinement was performed using each of the numerical methods to calculate interpolated map values and gradients: LI, QI and ID. For each refined model (from 100 × 4 × 6 × 3 = 7200 in total) we calculated the rmsd from the reference model, and then the average rmsds over the corresponding ensemble of 100 models. Figure 5 summarizes the results of this test. We observe that:

**Figure 5.**
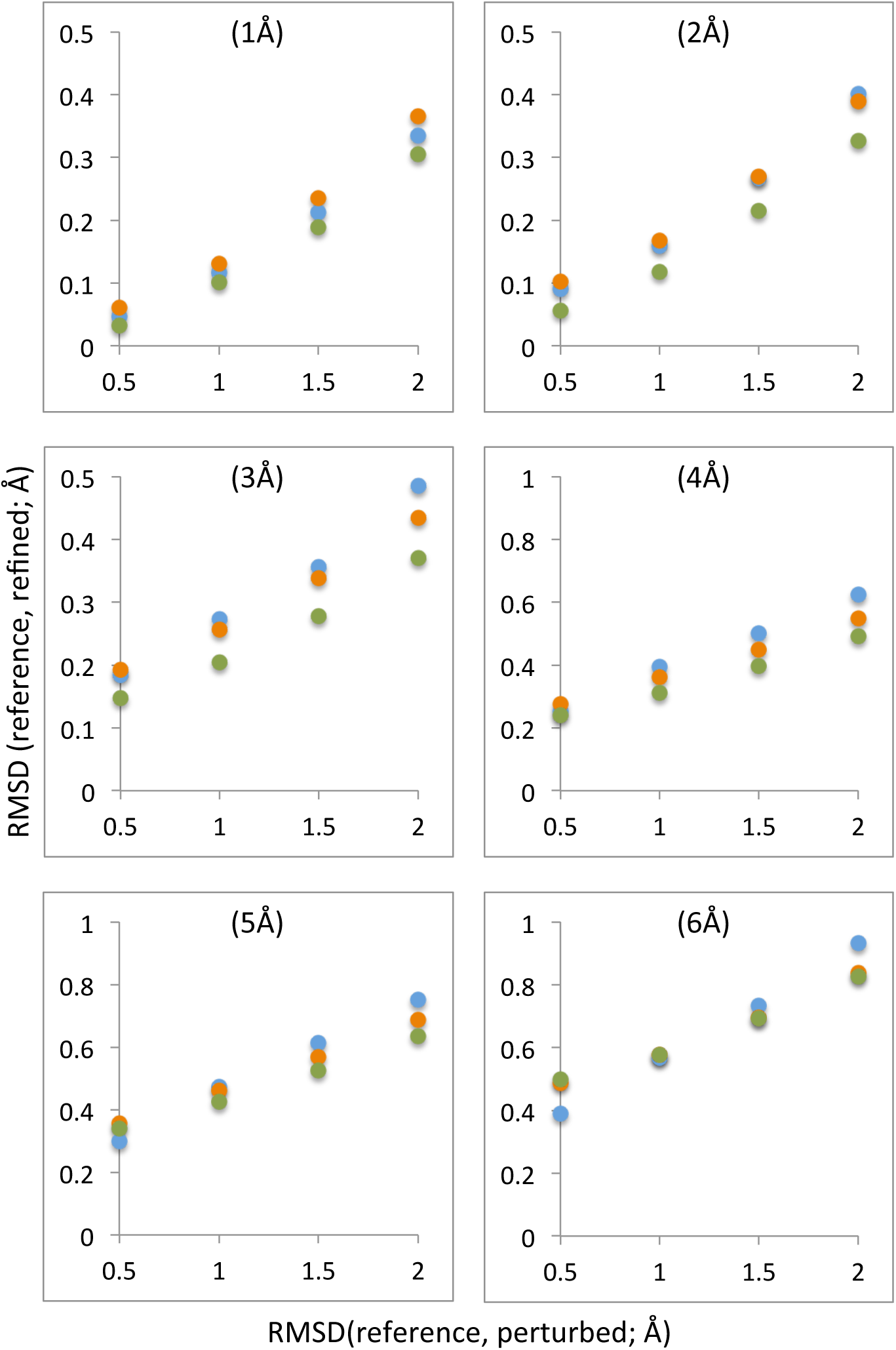
Refinement of perturbed models against maps of different resolutions using different interpolation methods: LI (orange), QI (blue) and ID (green). Horizontal axis shows rmsd between reference model and perturbed models: 0.5, 1.0, 1.5 and 2.0 Å. Vertical axis shows rmsd between reference model and refined models. Six plots correspond to six resolutions of the maps used (from 1 to 6 Å). See §3.1.2 for details.

- In most cases the difference between the results obtained using the three interpolation/gradient calculation choices is small compared to the magnitude of perturbation. Nevertheless, using the ID method produces systematically better results, agreeing with observations from refinements of the reference model. The QI approach gives slightly better results only at low resolution for models with relatively small starting errors. The difference in the calculation time between ID and other two methods is negligibly small. Consequently, ID is the default choice for *phenix.real_space_refine*.
- In all cases refinements were able to significantly reduce the difference between the reference and starting perturbed models. Refinement of models with the starting rmsd of 0.5 Å gives essentially the same results as refinement of non-perturbed reference model (similar rmsds). This shows the robustness of the approach.

To study how map sampling rate influences the results, we repeated the calculations from the previous test for the 3 Å resolution map calculated with a grid step of 0.2 Å (in previous tests it was *d_high_* /4 = 0.75 Å). We make two observations. First, using a finer grid nearly eliminates the differences among the three considered choices for interpolation/gradient calculations (LI, QI, ID). Second, the results obtained using ID are essentially the same as those using coarser maps. Both observations (corresponding plots are not shown) agree with results of similar tests using the reference model (see § 3.1.2).

To investigate the effect of map sharpness we considered four ensembles of perturbed models generated in the previous test. Each of these models was refined against a 3 Å resolution map calculated to correspond to different overall B values of the reference model, as described in § 3.1.2 (we did not repeat this test for other resolutions since it is computationally expensive). The results (not shown) confirm the same effect that we observed previously in tests with the reference model: an optimal map blurring or sharpening may result in a better refined model.

### 3.2 Refinement using data from PDB and EMDB

#### 3.2.1 Default refinement

In order to test the methods and demonstrate their utility we re-refined 385 cryo-EM models from the PDB that are reported at resolution 6 Å or better, that have model-map correlation greater than 0.3 and that contain only residues and ligands that are known to the *Phenix* restraint library. A number of metrics were analyzed: the model-to-map correlation coefficient CC_mask_ calculated in the map region around the model (see Afonine *et al*. (2018) for an exact definition), number of Ramachandran plot and rotamer outliers, excessive C_β_ deviations, the MolProbity clashscore (Chen *et al*., 2010) and EMRinger score (Barad et al., 2015; calculated for 277 entries with maps at 4.5 Å or better), all calculated for the initial models from PDB and for the models after refinement. Default parameters were used in refinement. The program ran successfully, generating a refined model for all cases, highlighting the robustness of the algorithms and the implementation. In all cases we observe substantial overall improvement of geometry metrics, such as reduced or fully eliminated Ramachandran plot and rotamer outliers, C_β_ deviations and MolProbity clashscore, as well as improvement of model to data (map) fit (Figure 6). The overall average EMringer score for initial models is 1.73 and for refined models is 2.26. The improvement of the EMRinger score for refined models indicates that amino-acid side chains are more chemically realistic and better fit the density map. Detailed validation or analysis of individual refinement results is outside the scope of this work.

**Figure 6.**
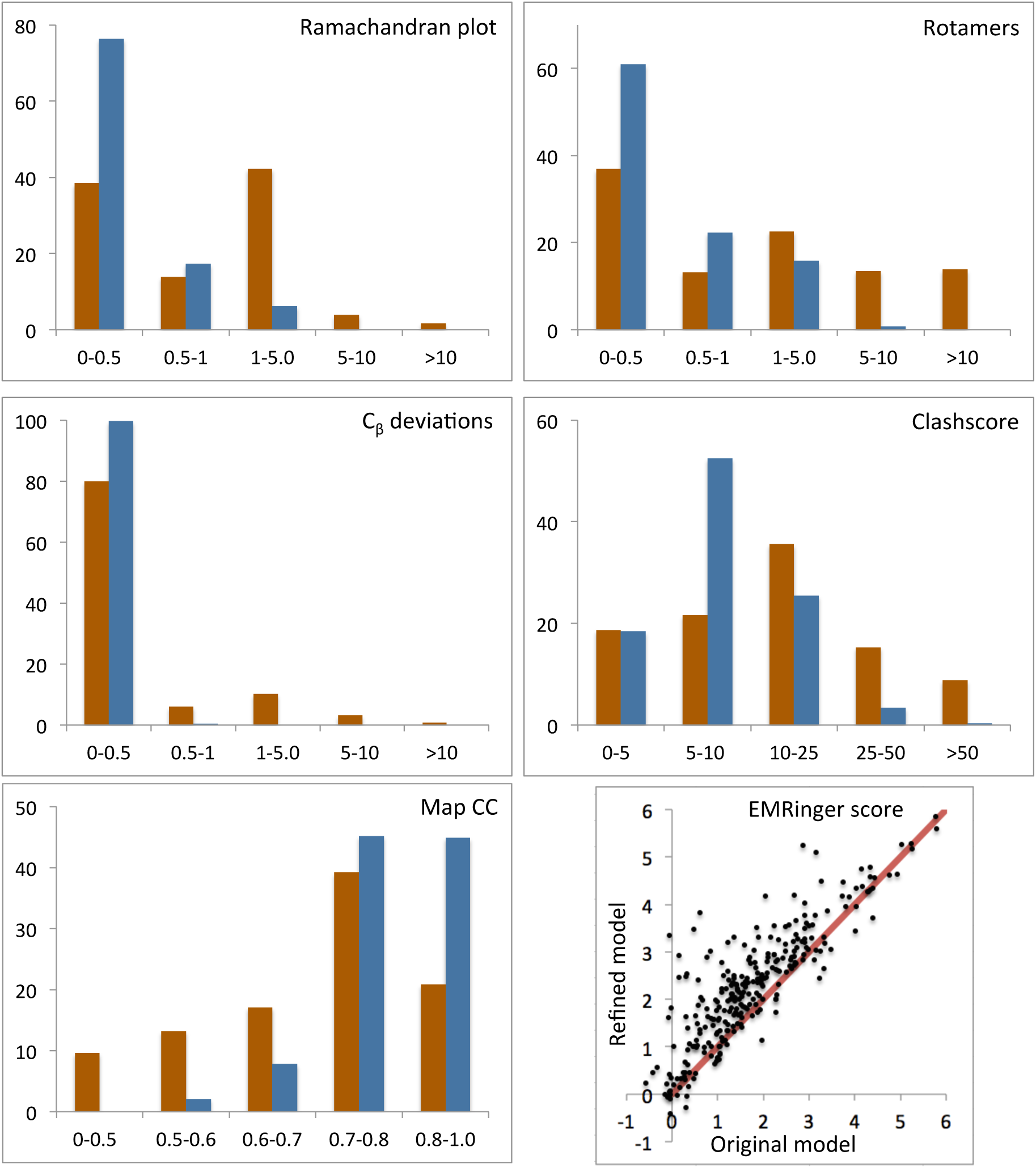
Model statistics before (brown) and after (blue) refinement using *phenix.real_space_refine*, showing Ramachandran plot and residue side chain rotamer outliers, C_β_ deviations, MolProbity clashscore and model-map correlation coefficient (CC_mask_). Scatter plot shows EMRinger score for original and refined models (resolution better than 4.5Å).

#### 3.2.2 Refinement against sharpened maps

Our tests using simulated data (§ 3.1) have indicated that map sharpening or blurring may be useful in refinement. To investigate this with the real experimental data we performed the following test. We selected models similarly to § 3.2.1, additionally requiring that independent half-maps had also been deposited by the researcher. This resulted in 76 entries. We performed test refinements against the first of the two half-maps and evaluated the refined model-to-data fit using the original second half-map that had not been used in any calculations. In two independent refinements, the first half map was taken either as deposited or modified with *phenix.auto_sharpen* (Terwilliger *et al*., 2018, submitted) to automatically optimally sharpen or blur the map. Figure 7 shows the model-map correlation CC_mask_ for models refined against the original and sharpened 1^st^ half-maps; the original 2^nd^ half-maps were used to compute the correlations. Overall, the CCs across all 76 cases are similar for refinement against the original 1st half-map and the sharpened 1*st* halfmap. Refinement models fit slightly but systematically better when using sharpened maps if the original model-map CC is low (<0.5) and systematically slightly worse if the original model-map is higher (CC>0.5). This agrees with the observation that target (7) allows for the removal of large errors but may slightly distort exact models (§3.1.2). Also, we note that MolProbity scores for models refined against sharpened maps are systematically better, but the difference is small.

**Figure 7.**
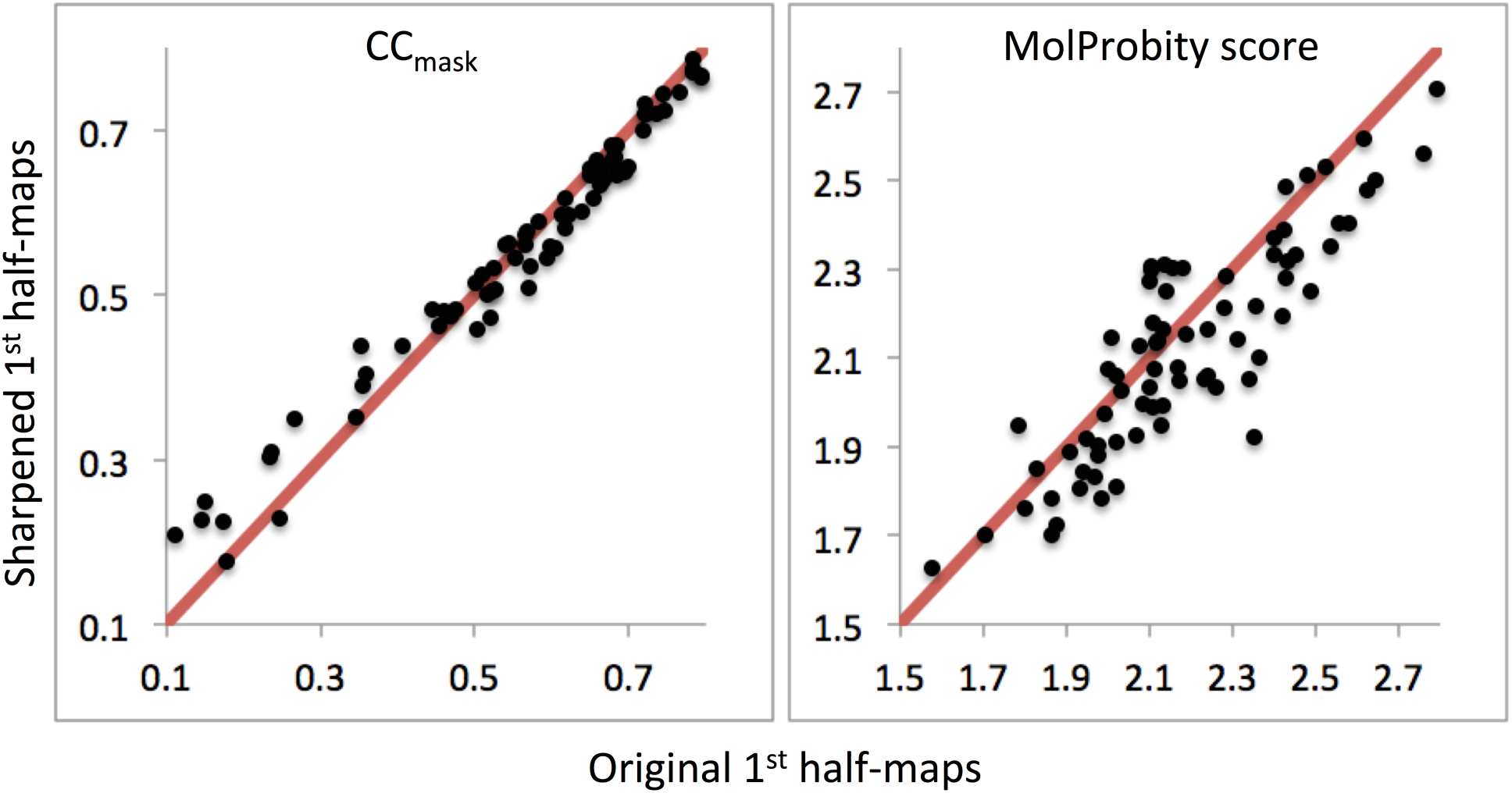
Left: correlation coefficient CC_mask_ calculated using original 2^nd^ half-maps and maps calculated maps from models refined against 1^st^ half-maps, original (x-axis) versus sharpened (y-axis). Right: MolProbity scores for models using original 1^st^ half-map versus sharpened 1^st^ half-map.

#### 3.2.3 Re-refinement of TRPV1 structure

The structure of the TRPV1 ion channel (PDB code 3j5p) was determined by single particle cryo-EM (Liao *et al*., 2013) at a resolution of 3.28 Å. The model was built manually and was not subject to refinement. As the model was not refined it contains substantial geometry violations: the clashscore is high (~100) and about one third of side chains are identified as rotamer outliers (Table 1). More recently, the better-resolved part of this structure has been re-evaluated using the same data (Barad *et al*., 2015; PDB code 3j9j, ankyrin domain not included). This involved some rebuilding and refinement using algorithms implemented in the Rosetta suite (DiMaio *et al*., 2015). The resulting model has much improved clashscore and EMRinger score (Barad *et al*., 2015) and no rotamer outliers, yet the number of Ramachandran plot outliers has increased compared to the original model (Table 1). We performed a refinement of 3j5p (the portion that matches 3j9j) using *phenix.real_space_refine* with all default settings and automatically, with no manual intervention, using the original, deposited map. The refinement took about 3 minutes on a Macintosh laptop. The refined model is similar to 3j9j (no rotamer outliers, much improved clashscore) but it also has no Ramachandran plot outliers, the EMRinger score is improved further and the model-to-map correlation (CC_mask_) is increased compared to both 3j5p and 3j9j.

## 4. Conclusions

Refinement of an atomic model against a map is increasingly important as the technique of cryo-EM rapidly develops. We have described the algorithms in a new *Phenix* tool, *phenix.real_space_refine*, that was specifically designed to perform such real-space refinements. While this work was inspired by rapid advances in the field of cryo-EM and the increasing number of three-dimensional reconstructions that allow atomic models to be refined (as opposed to rigid-body docked), the implementation is not limited to cryo-EM and crystallographic maps can be used as well. RSR is a natural choice for cryo-EM unlike crystallography, where real-space methods are complementary to Fourier-space refinement and are somewhat limited since crystallographic maps are almost always model biased.

The proposed real-space refinement procedure is fast due to using an atom-centered refinement target function that has been shown to be efficient at all tested resolutions, from 1 to 6 Å. Several options for key calculations steps, such as map interpolation, gradient calculation and preliminary processing of the target (experimental) map are available with the default choices selected on the basis of extensive test calculations. The real-space refinement algorithm includes a fast and efficient search for the optimal relative weight of restraints, a procedure that is extremely challenging for reciprocal-space refinement. The refinement algorithm is robust, with no failure for any of the cryo-EM tested. For all test model refinements improvements are observed; in some cases these improvements are significant. Future developments of the algorithms will include methods to account for local variation in map resolution, and efficient modelling of atomic displacements.

## Acknowledgement

This work was supported by the NIH (grant GM063210 to PDA, RJR and TT) and the *Phenix* Industrial Consortium. This work was supported in part by the US Department of Energy under Contract No. DE-AC02-05CH11231. AU acknowledges the support and the use of resources of the French Infrastructure for Integrated Structural Biology FRISBI ANR-10- INBS-05 and of Instruct-ERIC. RJR is supported by a Principal Research Fellowship funded by the Wellcome Trust (Grant 082961/ Z/07/Z).

## Appendix A. Real-space targets and convolution

We show here that, if the atoms all have the same shape, sampling a map at the positions of atomic centers, as in (7), can be made equivalent to the correlation function obtained by integrating or summing over the product of calculated and target densities, as in (3) or (5). Consider a simplified structure composed of a single atom. Looking for its best position according to (3) or (5) corresponds to seeking the position where the weighted average of the target map values (weighted by the atomic shape) inside a sphere centered at the trial atomic position is maximal. This calculation and check for the maximal value could be performed point by point. Alternatively, one can first calculate such averages forall grid points, replace the initial map values by these sums, and then simply choose the maximum. From a mathematical point of view this averaging can be considered as a convolution and, if calculated simultaneously for the whole map, can be performed rapidly (Lunin, in Urzhumtsev, 1985; Leslie, 1987; Urzhumtsev *et al.*, 1989). Checking the values of the averaged, i.e. blurred map, for their maximum corresponds to using targets (7) or (8). Below, we give a formal interpretation of these real-space targets.

Let *Z*_0_*f*_0_(|**s**|;*B*_0_) be a scattering factor of some isotropic atom characterized by a *B*_0_ value and the electron number Z_0_. Let *Z*_0_*ρ*_0_(**r**;B_0_) be an image of this atom in the corresponding model map if it is placed at the origin. Both *Z*_0_*f*_0_(|**s**|;*B*_0_) and *Z*_0_*#x03C1;*_0_(**r**;*B*_0_) are spherically symmetric and related by Fourier transformation. If a hypothetical structure is composed from a single atom positioned at **r**_0_, the corresponding model map is

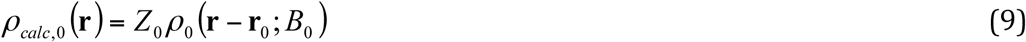

which can be seen as a convolution of a point scatterer at position r0 with the atomic shape. Due to the spherical symmetry of *ρ*_0_(**r**;*B*_0_), target function (3)

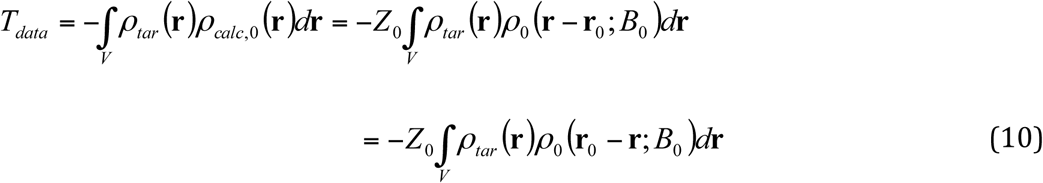

can be interpreted as a convolution of the target map with *ρ*_0_(**r**;*B*_0_) taken in point **r**_0_. Let {**F**_*tar*_(**s**)} be the set of Fourier coefficients corresponding to the target map *ρ*_*tar*_(**r**). By the convolution theorem, (10) is equal to the Fourier series of the corresponding Fourier coefficients:

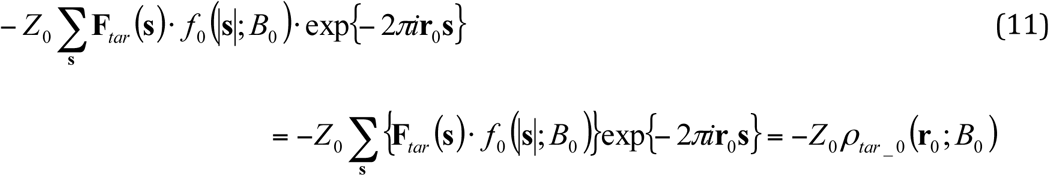

Here the map *ρ*_*tar*_0(**r**; *B*_0_) is a Fourier series calculated with the coefficients **F**_tar_(**s**)*f*_0_(|**s**|;*B*_0_).In other words, instead of blurring the model map with the atomic shape and calculating the point-by-point product of the two maps one may blur the experimental map and leave the model map unblurred, i.e. as a point map.

For a multi-atom model

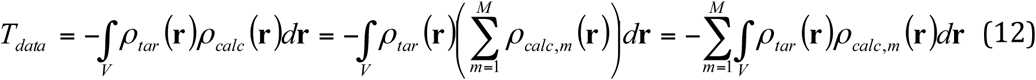

At resolutions typical for bio-crystallography the shapes of macromolecular atoms are similar. If we additionally suppose that all atoms of the structure have the same (or similar) atomic displacement parameters *B*_m_ = *B*_0_, then

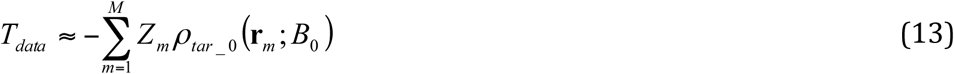

using the function *ρ*_tar_0_(**r**; *B*0) calculated once in advance. This shows that in calculating (8) we in fact implicitly sharpen the target map using *ρ*_tar_(**r**) instead of *ρ*_tar_0_(**r**; *B*0). Even when using (8) as the target, it is likely to be beneficial to choose an optimal sharpening factor, just as the signal in density correlations can be improved.

If the difference in atomic B values cannot be neglected, one can calculate in advance a few maps 
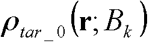
 for a range of B factor values *B_k_, k = 1, …, K*, and use the appropriate 
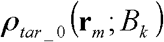
 for a particular atom *m*:

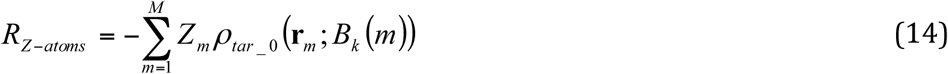

## Appendix B. 3D interpolation

Using atom-centered targets (7) and (8) requires an efficient and accurate interpolation of the maps calculated in three-dimensional regular grids. Several options have been considered in this work. Other choices for the interpolation (not considered in this work) are available (e.g. Lekien & Marsden, 2005) and their utility in the context of real-space refinement may be investigated in future works.

### B1. Trilinear interpolation

Consider a function *f*(*x*) of one variable in the interval (0,1). Let its values be known at the extremity of the interval, *f*_0_ = *f*(0) and *f*_1_ = *f*(l). A linear interpolation of this function is described as

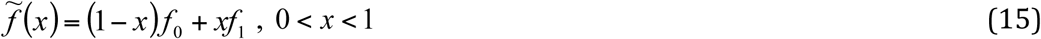

Exactly the same approach, known as a trilinear interpolation (https://en.wikipedia.org/wiki/Trilinearinterpolation), can be applied to constructing interpolations for the functions of three variables as described below.

If the values of the function in the corners of a unit cube are known, with notation *f*_000_,*f*_100_,*f*_010_,*f*_110_,*f*_001_,*f*_101_,*f*_011_,*f*_111_, then the interpolation formula is

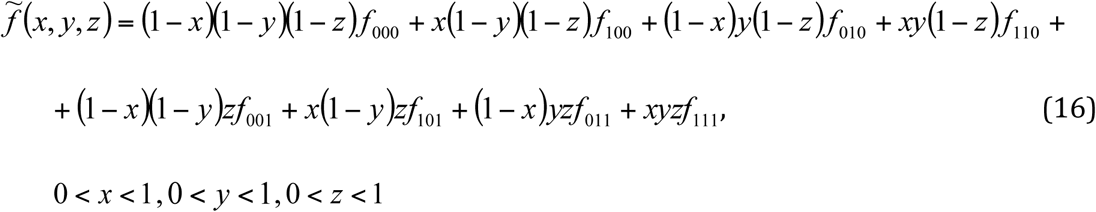

where each corner value is weighted by the volume of the parallelepiped opposite to this corner. For practical use we calculate it in three steps introducing first the intermediate interpolations

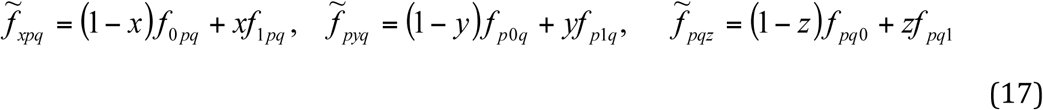

where *p* and *q* are either 0 or 1, then

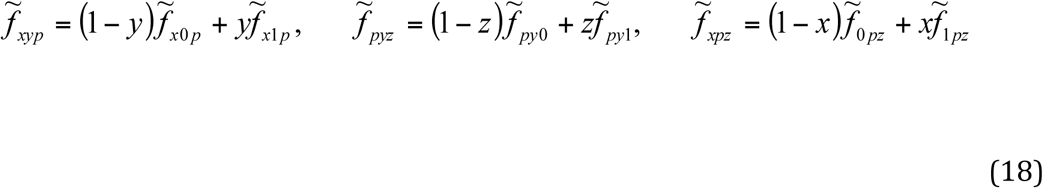

and finally

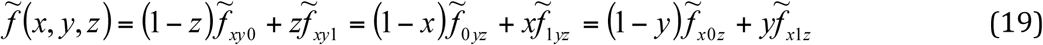

Expression (16) confirms that the result is independent of the order of intermediate interpolations in (17)–(19). This last formula gives the expressions for the partial derivatives of the interpolated function as

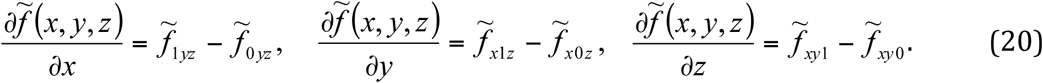

Now let a function *f*(*x,y,z*) be defined in fractional coordinates, on a grid with the step 
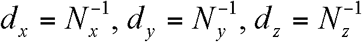
. Let’s consider a point (*x*_g_, *y*_g_, *z*_g_) and a ‘box’ of this grid that this point belongs to:

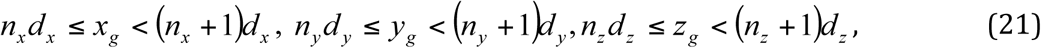

*n*_*x*_, *n*_*y*_, *n*_*z*_ being integer numbers. If we want to apply the interpolation formulae shown above, first we introduce intermediate variables rescaling this ‘box’ to a unit one:

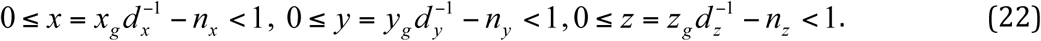

Resuming the whole algorithm, for each point (*x*_g_, *y*_g_, *z*_g_) where interpolation is required from the function defined on a grid with the steps *d*_x_, *d*_y_, *d*_z_, one

a. determines the grid box (21) which this point belongs to; e.g. if (*x*_g_, *y*_g_, *z*_g_) all are positive, *n*_x_, *n*_y_, *n*_z_ define the box corner closest to the origin
b. calculates new local interpolation coordinates (22) which can be considered as fractional with respect to the interpolation ‘box’
c. calculates *x*- and *y*-interpolated values (*y*-interpolated values are required only to calculate the derivatives)

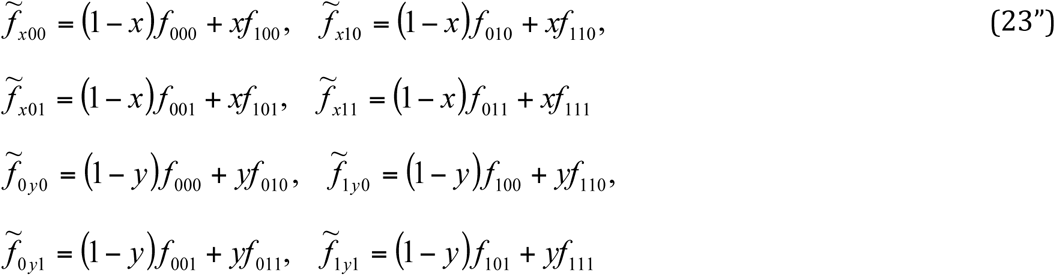
d. calculates *xy*-, *xz*- and *yz*-interpolated values (*yz*-interpolated values are required only to calculate the derivatives)

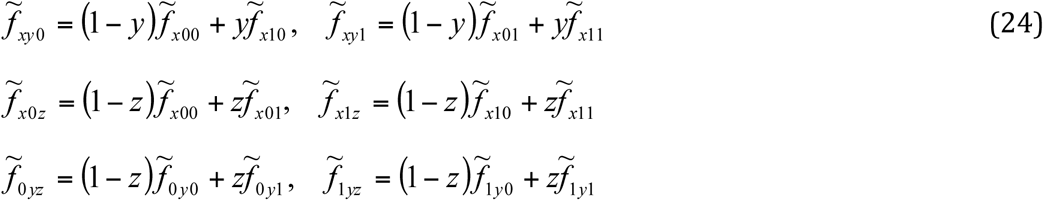
e. finally calculates *xyz*-interpolated value 

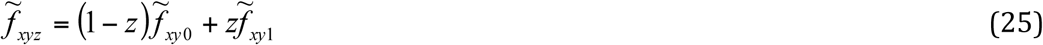

(considering (19), similar formulae could be used making the last interpolation by *x* or by *y*; the result is independent of this arbitrary choice).

### B2. Gradient calculation fo r the triple-linear interpolation

Trilinear interpolation means a linear interpolation in each of the three directions along the coordinate axes. As a consequence, straightforward estimates of the partial derivatives with respect to the initial coordinates are constants inside each interval (Figure 8a):

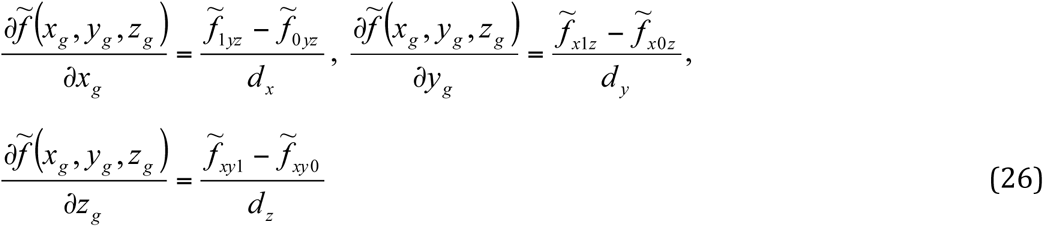

We will refer to this way of gradient calculation as LI, linear interpolation. One may note that in general the linear approximation that passes through the end points gives more accurate results for the function near these points, i.e. for (*x_g_*, *y_g_, z_g_*) close to the grid nodes, and less accurate at the middle of the interval. For derivatives estimated through (26) it is the opposite (see for example Figure 8a), and the derivatives are discontinuous as the point being evaluated moves from one box to its neighbor. Thus, a better estimate for the derivative of the original function sampled on the grid could be obtained if the point is in the middle of an interpolation interval (this gives an exact value for a quadratic function). To do so, we artificially construct a new interval around the point *x* in which we need to estimate the function derivative; the extremities of this new interval should be outside of the (0,1) otherwise the derivative will coincide with (26).

**Figure 8.**
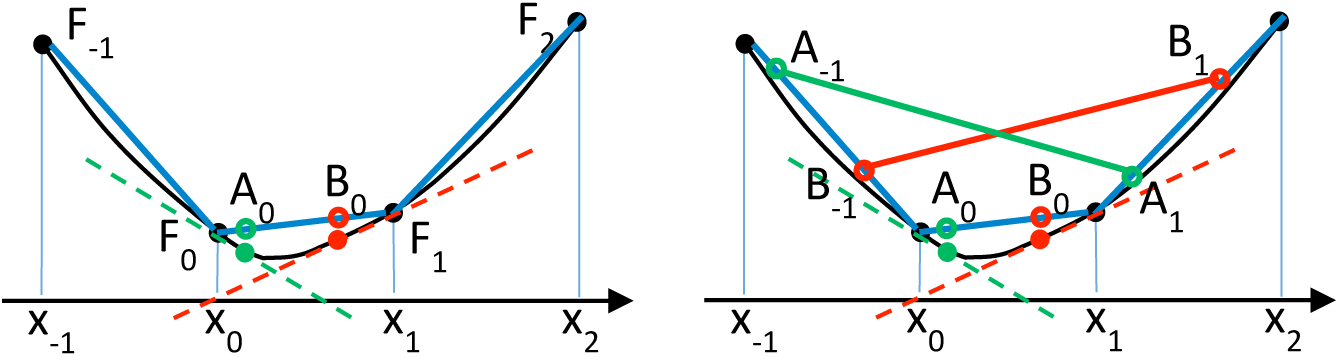
a) Black line shows a function *f*(*x*) whose values *F*_−1_, *F*_0_, *F*_1_ and *F*_2_ are known in equidistant (by *d*_*x*_) points *x*_−1_, *x*_0_, *x*_1_ and *x*_2_, respectively (black solid circles). Solid green and red circles show the true function values (considered as unknown) in the points inside the interval (*x*_0_, *x*_1_) in which we calculate interpolation for the function and its derivative. The slope of the dashed lines corresponds to the true derivative values at these points (also unknown). Blue lines illustrate the linear interpolation in the three intervals. Open circles in A_0_ and B_0_ illustrate the interpolated function values. The slope of the straight line (F_0_,F_1_) corresponds to the estimate (25) of the derivative *f*'(*x*) in points A_0_ and B_0_, the same value for both. Note that for point A_0_ even the sign of the estimate is wrong, b) Open circles in A_−1_, B_−1_, A_1_, B_1_ illustrate the interpolated function values in the points shifted by ± *d_x_* by the argument from A_0_ and B_0_, respectively. The slope of continuous green and red lines, corresponding to the derivative estimates using (26), is closer to that of the respective dashed lines.

To do so, for a point 0 ≤ *x* < 1:

- We define *x*_−1_ = *x* − 1 and *x*1 = *x* + 1; now *x* is in the middle of the interval (*x*−1, *x*_−1_); note that −1 ≤ *x*_-1_ ; 0 and 1 ≤ *x*1 ; 2; both here and below we suppose that the interval (0,1) is an internal one and there are grid points on both sides;
- We build linear interpolations at the intervals (−1,0) and (1,2) using *f*(−1) and *f*(0), and *f*(1) and *f*(2), respectively;
- Using these interpolations, we estimate function values
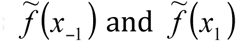
 for *x*_−1_ and for *x*_1_, respectively;
- We estimate the derivative in x using these values as (Figure 8b): 

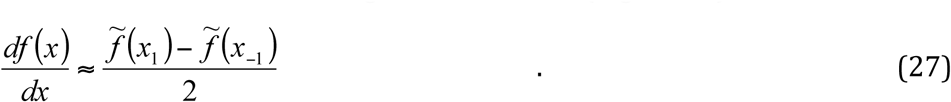

We expect to obtain more accurate results since in this method we use information from two intervals neighboring to that in which the derivative is calculated. For this reason we will refer to this way of gradient calculation as ID, interpolated difference or neighborinterval interpolation. Note that this approach gives the exact derivative value for quadratic functions.

For the initial three-dimensional problem, expressions, obtained by interpolation in three consecutive intervals by each variable, more accurate and preferable to (26), are

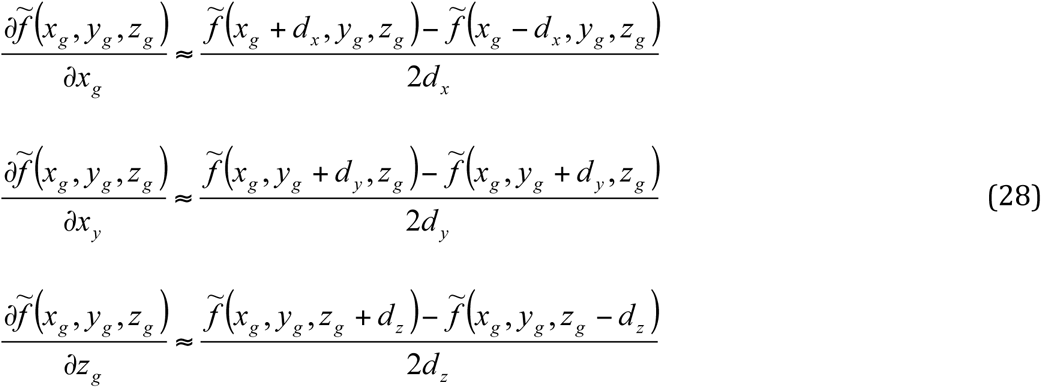

### B3. Quadratic interpolation

In the previous section, we discussed two procedures to calculate the gradient of a function defined on a grid By its nature, this function is continuous (because this is a density distribution), however we know its values only in the grid nodes, while off-nodes values are estimated by interpolation. As discussed above and illustrated by Figure 8, ID (28) may give a better than LI (26) approximation to the gradient of the *continuous function*. At the same time, expression (26) gives the exact gradient value of the *interpolated* function (16). Since it is the *interpolated and not the continuous function* that is actually used in minimization of(7), ID is less consistent with the tri-linear interpolation (16) than LI. Also, with ID we need to do three interpolations for each of three variables thus increasing the number of operations. Searching for a compromise, we introduced a quadratic interpolation (referred to as QI) for the function instead of the trilinear one.

As previously, we suppose that we know the values of a function *f*(*x,y,z*) at the integer points. The approximation we are looking for is

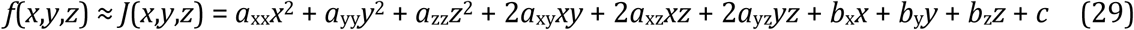

To determine the ten unknown coefficients we have eight values in the corners of the unit box, however only seven of them are independent since

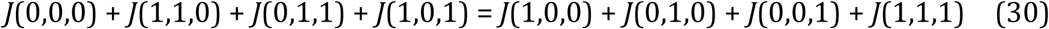

Three missing conditions can be obtained using function values at three other grid nodes outside the ‘box’ and closest to the point. For example, for 0 ≤ *x* < ½, 0 ≤ *y* <½, 0 ≤ z <, they are are *f*_(−1)00_, *f*_0(−1)0_ and *f*_00(−1)_ (as above,*f*_(−1)00_ stands for *f*(−1,0,0), etc); for ½ ≤ *x* < 1, ½ ≤ *y* < 1, ½ ≤ z < 1,they are values *f*_2_00, *f*_0_20 and *f*_002_, etc. The conditions (29)–(30) in the eleven selected grid points suggest the following procedure:

a. calculate the correction value 

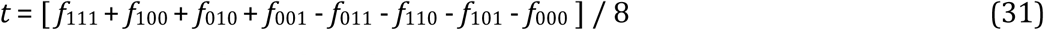
b. obtain the modified values that verify condition (30) 

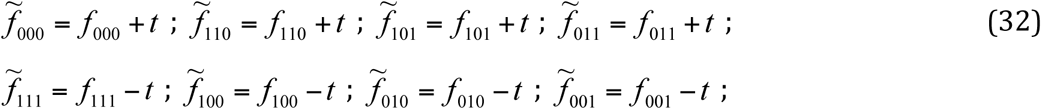
c. Calculate coefficients of the quadratic approximations 

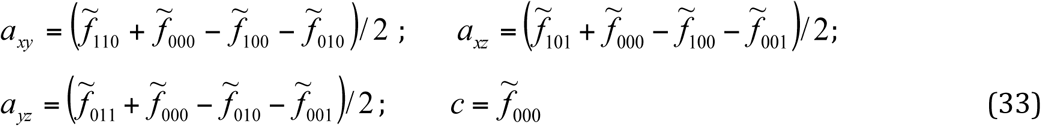

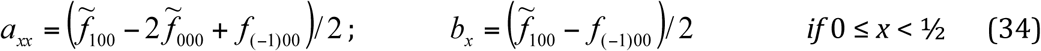

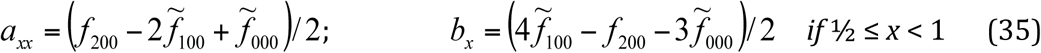

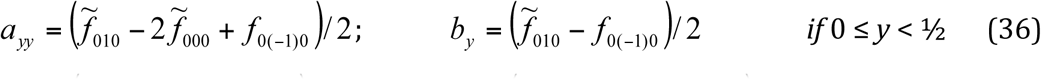

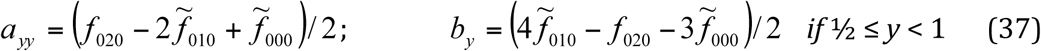

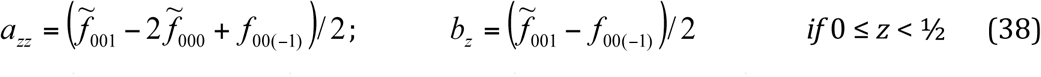

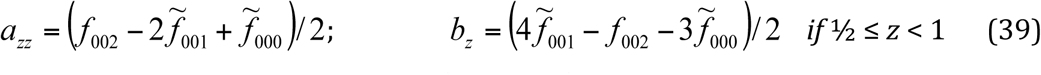
d. the interpolated function value

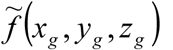
 is defined by (29) with the calculated coefficients and the partial derivatives are estimated, referring to the same formula, as

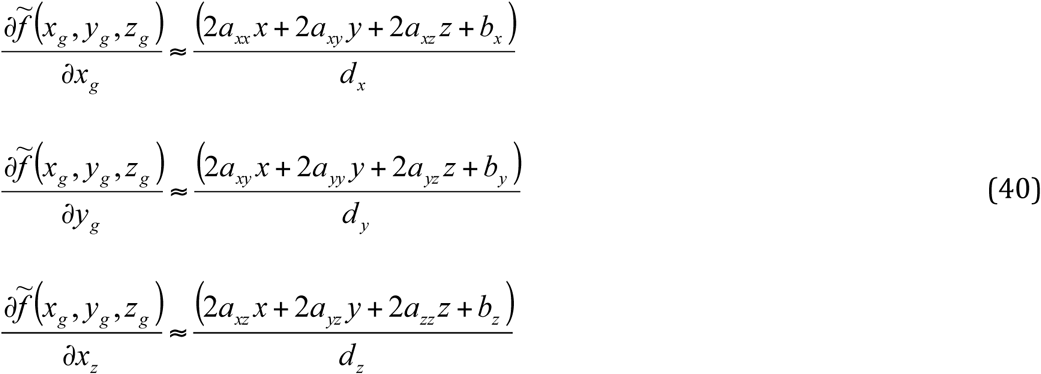

where *x,y, z* are defined through (*x_g_*, *V_g_*, *Z_g_*) as (21).

As a result of the advantages (faster calculation, consistency of the gradient calculation with the target function calculation) and disadvantages (less information used, required correction (32) of map values, general higher sensitivity of quadratic approximations to errors) of QI in comparison with ID, it may work better or worse depending on the circumstances, and § 3.1 compares the results of RSR using the three interpolation approaches discussed above.

## Footnotes

1 It is a widely known consequence of Parseval’s theorem (see for example, Diamond, 1971, or Arnold & Rossmann, 1988) that this is equivalent to a least-squares target between a full set of the corresponding complex Fourier coefficients; CNS (Brunger *et al.*, 1998) describes this as a “vector LS target”.

2 In crystallography, the set of the calculated Fourier coefficients usually coincides with that of the experimentally measured intensities.

